# Evolution of ACE2 and SARS-CoV-2 Interplay Across 247 Vertebrates

**DOI:** 10.1101/2021.01.28.428568

**Authors:** Tao Zhang, Qunfu Wu, Yicheng Ma, Wenjing Liu, Chengang Zhou, Zhigang Zhang

## Abstract

Severe acute respiratory syndrome coronavirus 2 (SARS-CoV-2) cause the most serious pandemics of Coronavirus Disease 2019 (COVID-19), which threatens human health and public safety. SARS-CoV-2 spike (S) protein uses angiotensin-converting enzyme 2 (ACE2) as recognized receptor for its entry into host cell that contributes to the infection of SARS-CoV-2 to hosts. Using computational modeling approach, this study resolved the evolutionary pattern of bonding affinity of ACE2 in 247 jawed vertebrates to the spike (S) protein of SARS-CoV-2. First, high-or-low binding affinity phenotype divergence of ACE2 to the S protein of SARS-CoV-2 has appeared in two ancient species of jawed vertebrates, *Scyliorhinus torazame* (low-affinity, Chondrichthyes) and *Latimeria chalumnae* (high-affinity, Coelacanthimorpha). Second, multiple independent affinity divergence events recur in fishes, amphibians-reptiles, birds, and mammals. Third, high affinity phenotypes go up in mammals, possibly implying the rapid expansion of mammals might accelerate the evolution of coronaviruses. Fourth, we found natural mutations at eight amino acid sites of ACE2 can determine most of phenotype divergences of bonding affinity in 247 vertebrates and resolved their related structural basis. Moreover, we also identified high-affinity or low-affinity-associated concomitant mutation group.The group linked to extremely high affinity may provide novel potentials for the development of human recombinant soluble ACE2 (hrsACE2) in treating patients with COVID-19 or for constructing genetically modified SARS-CoV-2 infection models promoting vaccines studies. These findings would offer potential benefits for the treatment and prevention of SARS-CoV-2.

## Introduction

An ongoing global pandemic of coronavirus disease 2019 (COVID-19), caused by the severe acute respiratory syndrome coronavirus 2 (SARS-CoV-2), have resulted in more than 70 million confirmed cases in 190 countries and more than 1.5 million deaths. The SARS-CoV-2 is a positive-strand RNA virus that causes severe respiratory syndrome in human. The genome of SARS-CoV-2 shares about 96% identity to the Bat Coronavirus BatCoV RaTG13[1]. SARS-CoV-2-like CoV was found in the pangolin species (*Malayan pangolins*), showing 91.02% identical to SARS-CoV-2[2]. Accordingly, *Rhinolophus affinis* and Malayan pangolin (*Manis javanica*) are considered as potentially natural hosts of SARS-CoV2[1, 2]. Aside from bat and pangolin, experiments with infectious SARS-CoV-2 suggested that SARS-CoV-2 replicates poorly in dogs, pigs, chickens, and ducks, but ferrets and cats are permissive to infection[3]. SARS-CoV-2 on mink farms was found to be transmitted between humans and mink and back to humans [4]. For non-human primates, *Macaca mulatta* is the most susceptible to SARS-CoV-2 infection, followed by *M. fascicularis* and *Callithrix jacchus*[5]. Syrian hamsters (*Mesocricetus auratus*) are also susceptible to SARS-CoV-2[6]. Anecdotal reports in a variety of news media reported that tigers in a New York Zoo tested positive for HCoV-19, in which these animals exhibited symptoms of the illness (https://www.nationalgeographic.com/animals/2020/04/tigercoronavirus-covid19-positive-test-bronx-zoo/). Thus, to resolving why the SARS-CoV-2 has both broad host ranges and various infection phenotypes is very important for the control of SARS-CoV-2. Some clues are implicated by two recent computer modeling studies[7, 8] but still remains limited.

ACE2 is the cellular receptor for SARS-CoV-2[3, 9-11]. Binding to ACE2 receptor is a critical initial step for the SARS-CoV-2 to enter into target cells[9]. Structural biologists have consecutively resolved structure of SARS-CoV-2 spike(S) protein and found SARS-CoV-2 S protein binds ACE2 with higher affinity than does SARS-CoV S protein[10], which may contribute to fast spread of COVID-19 from human to human, even human to animals (cat, tiger, and dog). Three following studies further provided deeply structural basis for the recognition of the SARS-CoV-2 by human ACE2 (hACE2), and found about 22 amino acid sites are involved in the interaction with the receptor binding domain (RBD) of spike protein of SARS-CoV-2[3, 11, 12]. ACE2 variants underlined interindividual variability and susceptibility to COVID-19 in Italian population[13]. Hence, it is very vital to elucidate whether natural and functional variations of the ACE2 determine both broad host ranges and diversified infection phenotypes of SARS-CoV-2 as well as to choose animal models, track down intermediate hosts, and develop recombinant or soluble ACE2 for the treatment of COVID-19. Even it would be helpful to understand why the bat species are natural reservoirs of SARS-CoV2 or SARS-CoV or SARSr CoV.

## Results

### Multiple independent affinity divergences between SARS-CoV-2 and ACE2 in different lineages of 247 Vertebrates

We obtained 247 complete ACE2 protein sequences about the length of 800 amino acids from NCBI and Uniprot databases. Those ACE2 proteins represent 247 jawed vertebrates, belonging to Chondrichthyes, Coelacanthimorpha, Cladistia, Actinopteri, Amphibia, Crocodylia, Testudines, Lepidosauria, Aves and Mammalia (including *Homo species*) (**Supplementary Data Table S1**). All 247 ACE2 protein sequences were aligned and the regions that ranged from 19 to 619 amino acid sites referred to hACE2 were used to construct ACE2 protein tree and to perform homologous modeling of SARS-CoV2 RBD–ACE2 complex using Swiss-Modeling (https://swissmodel.expasy.org/). Binding affinity (1/*K*_d_) is typically measured and reported by the equilibrium dissociation constant (*K*_d_), which is used to evaluate molecular interactions[14]. The smaller the *K*_d_ value, the greater the binding affinity of the ligand for its target. The protein-protein binding affinity of ACE2 and SARS-CoV-2 S RBD were estimated using online PRODIGY tool[15, 16] based on swill-modeling results. To estimate binding affinity phenotypes of jawed vertebrate ACE2 and SARS-CoV2 RBD (**Figure 1, Figure S1 and Supplementary Data TableS1**), all estimated *K*_d_ values were normalized using the *K*_d_ value of hACE2 and S protein interplay for obtaining relative differential expression pattern of binding affinity as follows: *K*_d others_/ *K*_d homo_>1, defined as lower affinity than human; *K*_d others_/ *K*_d homo_=1, the same affinity to human; *K*_d homo_/ *K*_d others_ >1, higher affinity than human.

**Figure 1.**
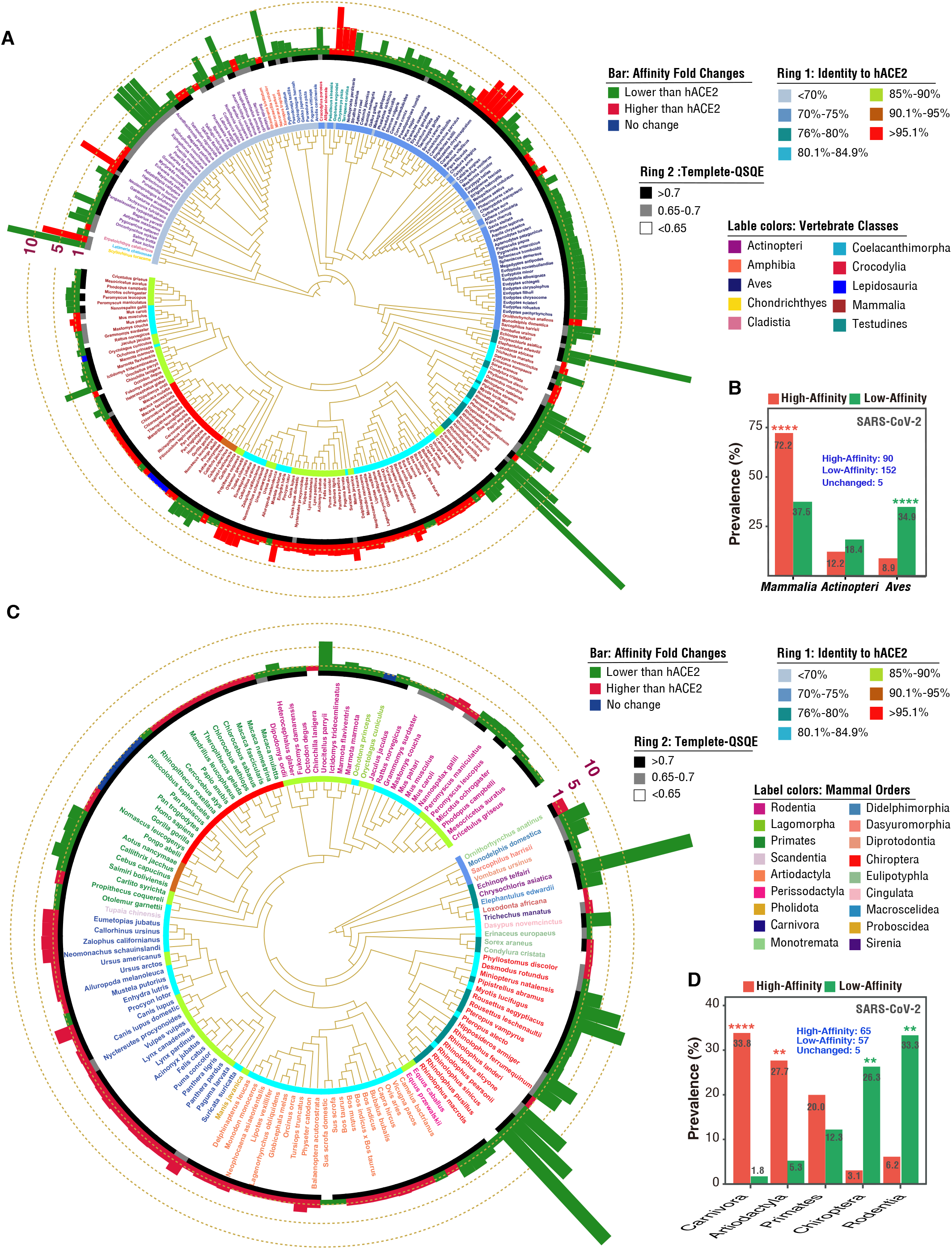
Predicted bonding affinities between ACE2 of 247 jawed vertebrates and the RBD of the S protein of SARS-COV-2. Linked to Figure S1. **A**. ACE2 protein tree 247 vertebrates with affinity fold change relative to hACE2. Red bars indicated higher affinity (1/*K*_d_) than hACE2. Green bars indicated lower affinity (1/*K*_d_) than hACE2. **B**. Enrichment analysis of affinity phenotypes in Mammalia, Actinopteri, and Aves using Fisher exact test under p-value <0.05 corrected by Benjamini-Hochberg (BH) method. **C**. ACE2 protein tree 127 mammals with affinity fold change relative to hACE2. Red bars indicated higher affinity (1/*K*_d_) than hACE2. Green bars indicated lower affinity (1/*K*_d_) than hACE2. **D**. Enrichment analysis of affinity phenotypes in Carnivora, Artiodactyla, Primates, Chiroptera, and Rodentia orders from mammals using Fisher exact test under p-value <0.05 corrected by the BH method.

We firstly revealed the divergence of high-or-low affinity between ACE2 and SARS-CoV2 occurs in two most ancient jawed vertebrates Chondrichthyes (about 16 times lower than human, *K*d(hACE2) = 2.1nM) and Coelacanthimorpha (nearly 10 times higher than human) (**Figure 1A and Figure S1A**). During the evolution of jawed vertebrates, we gradually unveiled multiple independent events of binding affinity divergence appearing in Actinopteri, Amphibia, Lepidosauria, Aves and Mammals. Compared with Mammals, high-affinity phenotypes are rare in Actinopteri (28%) and Aves (13%) (**Figure 1B, Figure S1B and Supplementary Data TableS1**). In mammalian species, the prevalence of high-affinity phenotypes is up to 51%, indicating the presence of mammal species might drive fast evolution of SARS-CoV-2 or SARS-CoV-2-like CoVs. We further found distinct binding affinity divergences in the different orders of mammals (**Figure 1C-1D and Figure S1C-1D**). High prevalence of high-affinity phenotypes appears in Artiodactyla (86%) and Carnivora (96%) (**Figure 1D and Figure S1D**). Different from Artiodactyla, Perissodactyla species (Horses) which is the most closely related to Artiodactyla show low-affinity phenotypes (7.3nM). Low-affinity phenotypes were dominant in Chiroptera (88%) and Rodentia (83%). Moreover, the *K*d values (14.56±4.08 nM) (mean±S.E.) of Chiroptera species are significantly higher than those (3.63±0.51 nM) of Rodentia species. *Rhinolophus sinicus* showed high *K*d value of 68nM. This finding suggests most of Chiroptera species have low affinity or high tolerance to SARS-CoV-2 and theoretically can be considered as the most suitable carriers of SARS-CoV-2 or SARS-CoV-2-like CoVs. In Primates (54% high and 29% low) and their closely outgroup Scandentia, we found slightly high or consistent affinity phenotypes in Old World Monkeys (OWMs) with one exception of Golden snub-nosed monkey with weakly low affinity. In contrast, low-affinity phenotypes appeared in New World Monkeys (NWMs), *Tarsiiformes, Lorisiformes* as well as Chinese tree shrew (Scandentia order as the closely relative of Primates).

Surprisingly, Coquerel’s sifaka (*Propithecus coquereli*) belonging to ancient primates (Lemuriformes species) shows high affinity (1.2nM). In other rare orders of mammals, we also found the presence of high-or-low affinity phenotypes, such as the most ancient mammal Platypus (*Ornithorhynchus anatinus*) showing high affinity (0.83nM) and Cape golden mole (*Chrysochloris asiatica*) showing greatly low affinity (52nM).

We systematically explored the ACE2 affinity to the S protein of SARS-CoV-2 across 247 vertebrate hosts. 54 animal species from our studied cohort were experimentally confirmed by published or pre-printed reports so far, including 48 susceptible and six uninfected animal species[3, 6, 17-25]. In this study, 39 of 48 reported infected animals had highly predicted affinity and 5 of 6 experimentally showing no infection were predicted having low affinity. Accordingly, high consistence between our predicated and experimentally confirmed phenotypes suggests the reliability of predicted phenotypes in this study.

### Detecting the effect of known amino acid sites in ACE2 involved in the interaction with SARS-CoV-2 on affinity divergences in vertebrates

To investigate what amino acid variations in ACE2 leading to affinity divergences between ACE2 and the S protein of SAR-CoV-2, we firstly characterized all mutations of 247 vertebrates in known 22 amino acid sites[12] determining the function of ACE2 binding to S protein and then estimated affinity changes of all mutants by mutating corresponding amino acids in hACE2. According to affinity fold changes of mutated type (MT) and wild type (WT), we found 55.9%, 76.5%, and 91.9% of WT can be recovered in MT below 1.41, 2.00, and 4.00-fold changes, respectively (**Figure S2**). These results suggested that the variations of known 22 amino acid sites were unable to completely explain our predicted affinity phenotype variations.

To determine what natural variations in ACE2 protein determine our predicted affinity divergence, we selected top 3 species with high-or-low affinity in Actinopteri, Aves, and Mammals and detected amino acid changes corresponding to high-or-low affinity phenotypes, which were shared by at least two species with consistent phenotype but absent in species with converse phenotype (**Figure 2A**). All detected amino acid variations were traced into two ancient species *Scyliorhinus torazame* (low-affinity, Chondrichthyes) and *Latimeria chalumnae* (high-affinity, Coelacanthimorpha) (Figure 2A). Total 64 amino acid sites were detected possibly involved in altering bonding affinity of ACE2 to S protein (**Figure 2A**), which showed host specificities of fishes, birds, and mammals. To confirm whether those amino acid changes in each species lead to similar phenotypes, we constructed amino acid mutants in hACE2 and tested bonding affinity of the mutants to S-protein. We found wild phenotypes can be almost recovered in hACE2 (**Figure S3**). Crossed amino acid replacements linked to high or low affinity phenotypes in hACE2 also confirmed phenotypic reversal (**Figure S3**). These results suggested our detected conserved amino acid changes had potentials to yield observed phenotype divergences. By testing affinities of different mutations, we found that natural variations of 16 amino acid sites might be linked to affinity divergences of top 3 species with high-or-low affinity in Actinopteri (e.g., Q27 and A27), Aves (e.g., N27 and I27), and Mammals (e.g., Q79 and H41) (**Figure 2B**). These findings suggested multiple independent amino acid mutation events might contribute to convergent affinity divergences in different jawed vertebrate lineages.

**Figure 2.**
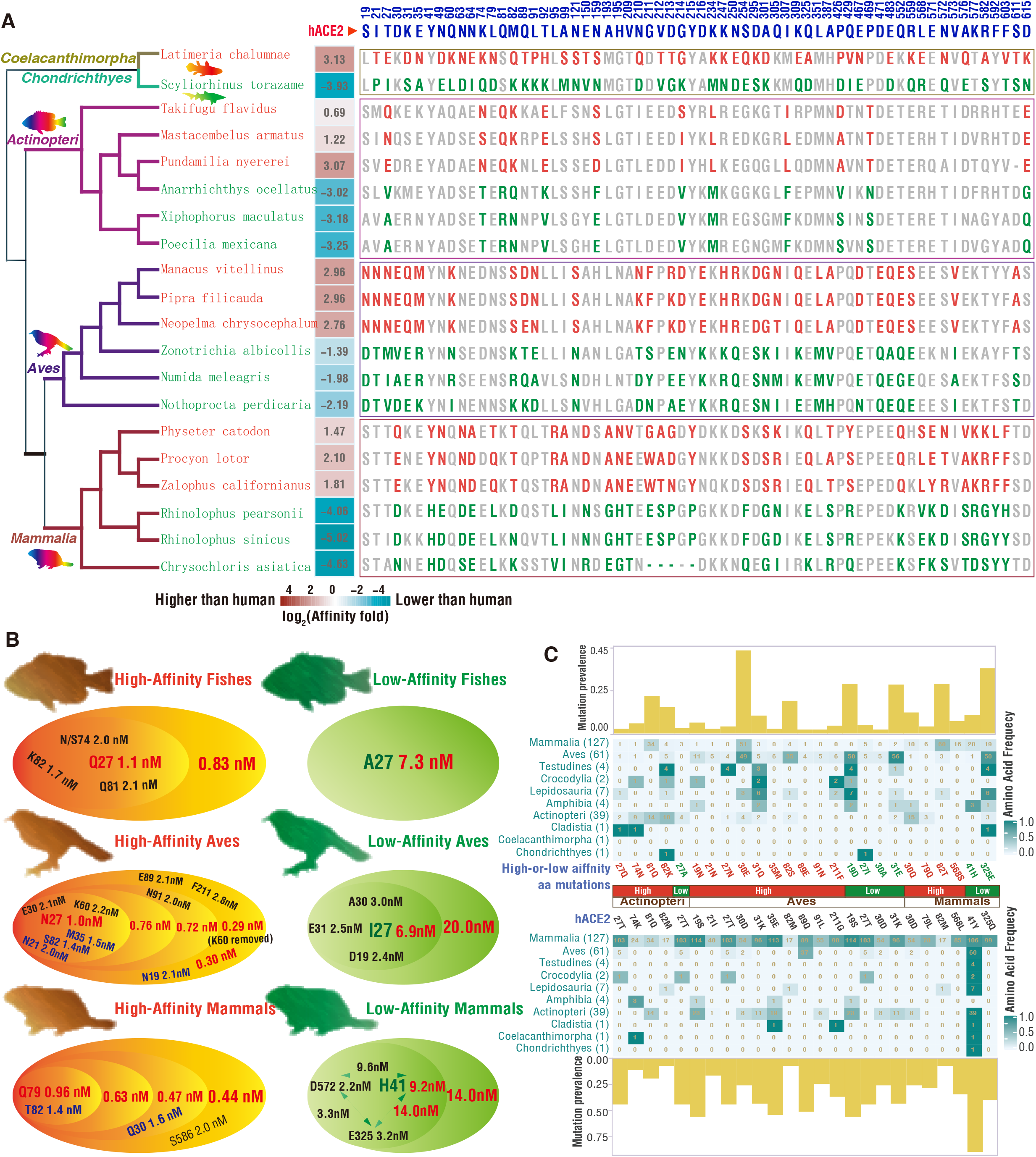
Amino acid changes corresponding to high-or-low affinity phenotypes from 247 vertebrates. Linked to Figure S3. **A**. Amino acid (AA) changes corresponding to high-or-low affinity phenotypes in top 3 high and 3 low affinity hosts from Actinopteri, Aves, Mammalia classes. High affinity AA changes were marked by red color, low affinity AA changes were marked by green color. High-or-low affinity associated AA changes were shared by at least two species with consistent phenotype but absent in reverse phenotype. For *Latimeria chalumnae* and *Scyliorhinus torazame*, affinity-associated AA changes were based on the AA combination of Actinopteri, Aves, Mammalia classes. Heatmap indicated the fold change in bonding affinity relative to hACE2 when replaced hACE2 with the affinity-associated AA changes from the corresponding non-human animals. Positive values in heatmap mean increase folds and negative values mean decreased folds relative to hACE2. **B**. Composition of AA changes leading to top high or low affinity combined with predicted affinities of mutated hACE2 in Actinopteri, Aves, and Mammalia classes. **C**. Frequency of specific amino acid changes linked to high-or-low affinities in Actinopteri, Aves, Mammalia classes across different vertebrate lineages.

To further determine what amino acid changes result in high-or-low affinity phenotype divergences in fish, bird, and mammals, we performed hACE2-based affinity tests using step-by-step single amino acid mutation of conserved amino acid residues associated with binding affinity changes in four vertebrate animal pairs. Our results showed that Q27 and A27 led to most of high-and-low affinity divergences in fishes (**Figure 2B and Figure S4**). N27 and I27 caused crucial differences of high-and-low affinity phenotypes in birds (**Figure 2B and Figure S5**). Q79 and H41 produced major divergences of high-and-low affinity in mammals (**Figure 2B and Figure S6**). Other amino acid mutations relying on those key amino acid mutations can strengthen or weaken bonding affinities (**Figure 2B, 2C**). Combined with phenotype testing based on step-by-step single amino acid mutation (**Figures S4-S7**), we speculated that concomitant amino acid mutations of 19-82 amino acid region (referring hACE2) might be involved in affinity divergences in jawed vertebrates. Even so, we found 41Y to H mutation only covering 20 of 41 low-affinity mammal species and five species in non-mammals (**Figure S8**), indicating that novel mutations still need to be further investigated.

### Novel amino acid mutations contributing to recurred affinity divergences in vertebrates

By performing the alignment of 19-82 amino acid region of 247 vertebrates, we found candidate amino acid changes of eight conserved amino acid sites including 19, 24, 27, 30, 34, 41, 79, and 82 (referred to as hACE2) associated with phenotype divergences (**Figures 2 and 3, Figure S8**). To confirm this observation, we conducted a series of affinity tests by mutating all single amino acids at each of the 8 sites from 247 vertebrates in hACE2 (**Figure 3A**). S19 and T27 potentially linked to low-affinity phenotype is dominant in mammals and D19 and E27 is in non-mammals. Q24 deletion leading to low affinity appear in non-mammals. Q30 and N30 resulting in high affinity appeared in both mammals and non-mammals. K30 and S30 could bring about high affinity in non-mammals. R34, K34, and Y34 (specific to Carnivora) may contribute to high affinity. 79M was mainly associated with low affinity in Artiodactyla order species of mammals. All other mutations at site 79 could be linked with high affinity. Mutations at site 82 also contribute to increased affinity. Totally, mutations with lower affinity at sites 24 and 34 could explain low-affinity phenotypes in about 50% mammal species lack of H41 (**Figure 3A**). By screening those mutations with the most extreme affinity at any one of eight sites, we preliminarily identified a high-affinity-associated concomitant mutation group (N(P)19-N27-Q30-Y41-Q79-T82) and low-affinity-associated concomitant mutation group (L19-24Del-A27-N34-H41-M79-M82) (**Figure 3B and 3C**). By performing consistent mutations in vertebrate species with converse phenotypes, we found candidate functional concomitant mutations obviously reversed bonding affinity phenotypes across vertebrates (boosted affinity in **Figure 3B** and reduced affinity in **Figure 3C**). These results further suggested that amino acid variations of eight conserved amino acid sites (referred to as hACE2: 19, 24, 27, 30, 34, 41, 79, and 82) across 247 jawed vertebrates might contribute to bonding affinity divergences with the SARS-CoV-2 S protein.

**Figure 3.**
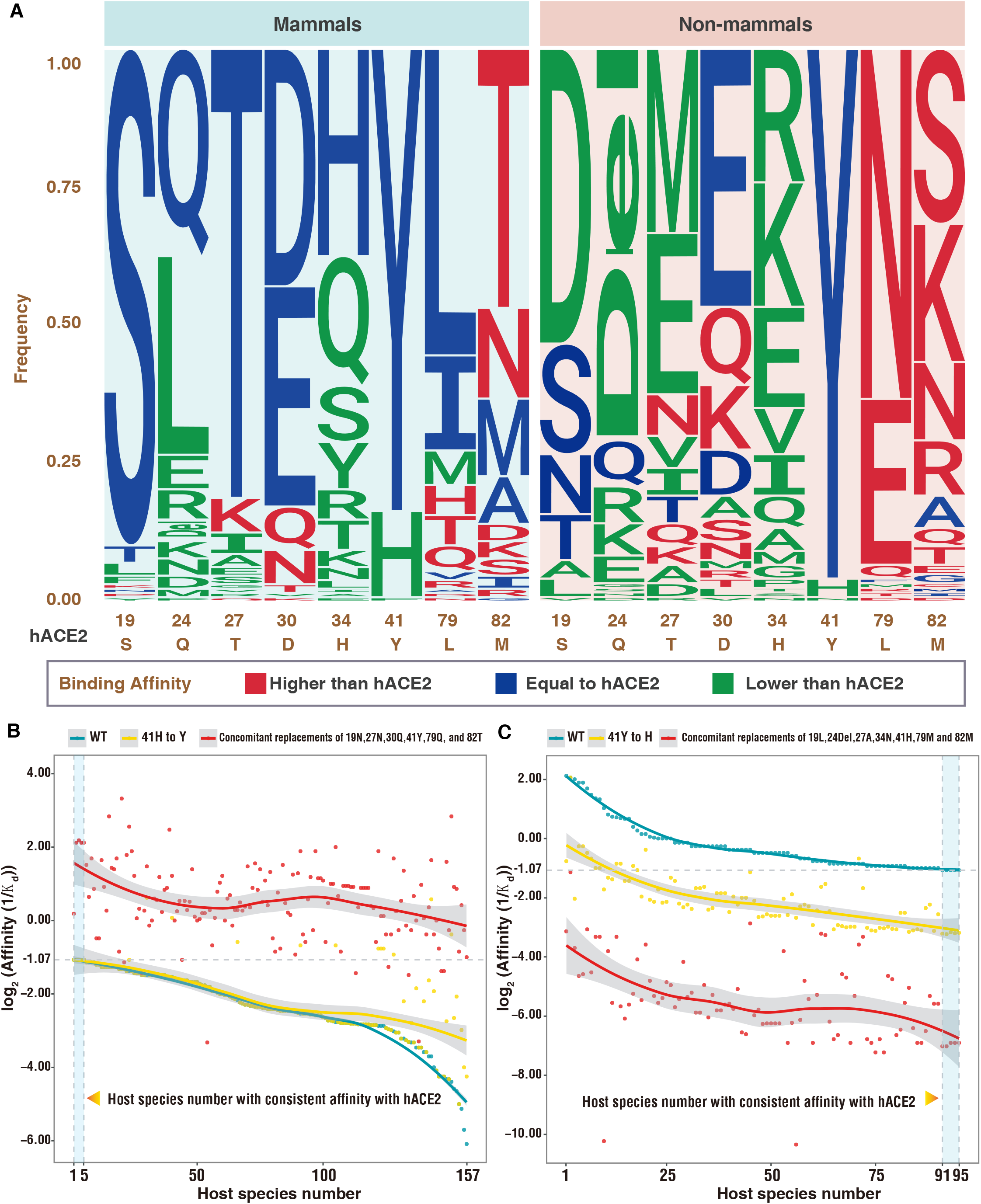
Amino acid characteristics at eight conserved amino acid sites leading to high-or-low affinity divergence in 247 vertebrates. Linked to Figures S4-S8. **A**. Frequency of amino acid residues at eight conserved amino acid sites in mammalian hosts and non-mammalian hosts contributing to different affinity phenotypes after mutated in hACE2 independently. Higher than wild hACE2 affinity marked by red; equal to hACE2 by blue; and lower than hACE2 by green. **B**. Affinity renversement of low-affinity animal hosts when related amino acids were replaced by high-affinity amino acid group (19N,27N,30Q,41Y,79Q, and 82T). The 1-5 vertebrates with affinity equal to wild hACE2 affinity and 6-157 vertebrates with affinity lower than wild hACE2 affinity. 41H to Y was used as control to eliminate the effect of 41H linked to extreme low affinity. **C**. Affinity renversement of high-affinity animal hosts when related amino acids were replaced by low affinity amino acid group (19L,24Del,27A,34N,41H,79M and 82M. The 1-90 vertebrates with affinity higher than wild hACE2 affinity and 91-95 vertebrates with affinity equal to wild hACE2 affinity. 41Y to H was used as control due to its contribution to extreme low affinity.

### Structural basis determining affinity divergence of different lineages in vertebrates

We found there are the mutations of eight amino acid sties in ACE2 contributing to significant SARS-CoV-2-interacting changes (**Figure 4**). The presence of polar interaction bonds or the shrink of contacting face contributed by amino acid mutations can increase bonding affinity of ACE2 to the S protein. For example, compared with hACE2, N27/Q27 residues form novel polar bonds to Y473 in the S protein of SARS-CoV-2, which can increase bonding affinity implicated by lower *K*d values (1.0nM and 1.1 nM, respectively). Q30 (1.6nM), Q79 (1.0nM) and T82 (1.4nM) showed higher affinity mainly due to the shrink of interacting face to S protein. In contrast, the loss of polar bonds or the enlargement of contacting face to S protein determine lower bonding affinity of ACE2 to S protein. For example, H41 lost hydrogen bond to both N501 and T500 of S protein and then resulted in the lowest bonding affinity (9.2nM). The deletion of Q24 also caused low affinity (4.3nM). By enlarging the contacting face with the S protein, the residues A27 and I27 showed the second low bonding affinity (7.3nM and 6.9 nM). L19, N34, and M79 weakly reduced bonding affinity by changing the interacting face between ACE2 and S protein. Those predicted structural interacting changes can potentially support extreme phenotype divergences of bonding affinity between ACE2 and the S protein of SARS-CoV-2.

**Figure 4.**
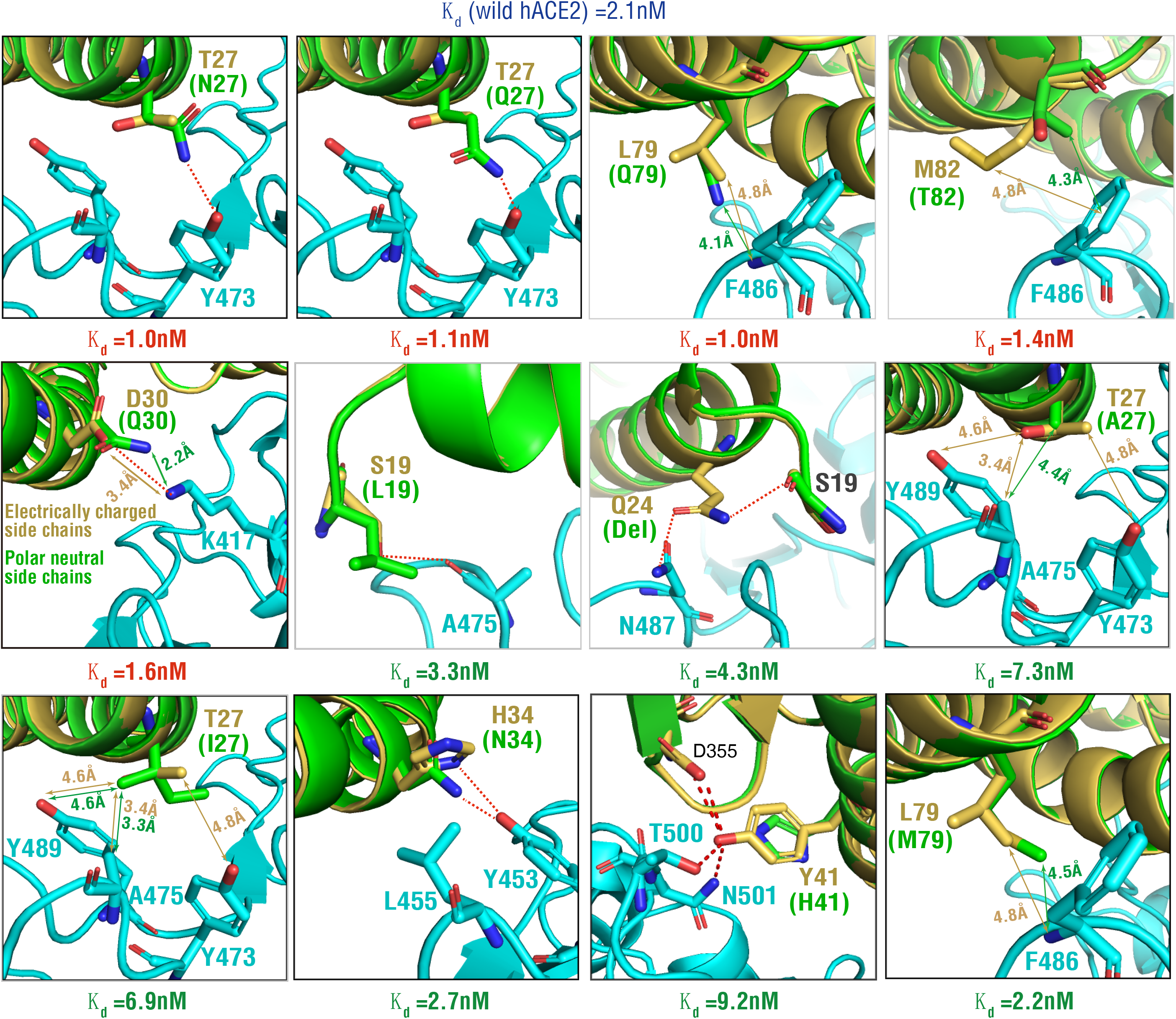
3D complex structure corresponding to key amino acid changes associated with affinity divergence. Linked to Figure 3. 3D complex structures were reconstructed based on those key amino acid changes resulting in high-or-low affinity divergence. Amino acids from wild hACE2 was marked by golden color. In brackets, high-or-low-affinity associated amino acid residues were marked by green. Amino acids and structures marked by blue color belonged to the RBD of the S protein of SARS-CoV-2.

## Discussion

Multiple studies have confirmed that ACE2 is the cellular receptor for SARS-CoV-2 and have found about 22 key amino acid sites of human ACE2 (hACE2) to be responsible for the interaction with SARS-CoV-2[1, 3, 9, 12]. However, whether natural ACE2 variants from various vertebrates could contribute to probable SARS-CoV-2 infections to non-human animals or transmissions from animals to humans is still unclear, although computational modeling, cell studies and animal experiments implicated that SARS-CoV-2 might infect non-human animals, such as some mammals including civet, ferrets, dog, cat, mink, pangolin, and so on. Our predicted affinity phenotype divergences recurring in different lineages (the oldest species, bony fishes, birds, and mammals) in 247 jawed vertebrates led to a possibility that the SARS-CoV-2 or SARS-CoV-2-like viruses are experiencing relaxed selection, which might partially contribute to intermittent appearances of diversified CoVs in recent years, such as SARS-CoV (2002-2003 years)[26], MERS-CoV (2012-2015 years)[26], and SARS-CoV-2 (2019 years to date).

Few high-affinity non-mammalian hosts, such as those species in *Cichlidae* family of Actinopteri, *Pipridae* family of Aves, and Testudines might have possible risks for SARS-CoV-2 infection. Nevertheless, considering those animals as potential intermediate hosts of SAR-CoV-2 could be ignored due to their habitats apart from human or small population sizes or potential effects of other unknown functional molecules. According to this perspective, Turtles were thought to be potential intermediate hosts[27] is unconvincing. For animals with low affinity phenotypes, the predicted affinity (2.8nM) of chicken is lower than hACE2 affinity (2.1nM), consistent with the report that chicken was not susceptible to SARS-CoV-2 infection[3, 28]. Our predicted low affinity of snakes (*Ophiophagus Hannah*, 5.8nM; *Pseudonaja textilis*, 6.9nM; *Python bivittatus*, 7.2nM) did not support that snakes were thought to be potential intermediate hosts[27]. On June 14 (2020), SARS-CoV-2 virus was detected in the cutting board of imported salmon (Chinook salmon) (*Oncorhynchus tshawytscha*) rising the discussion of whether salmon fish could be infected by SARS-CoV-2. Chinook salmon was not included in our studied cohorts but its close relative rainbow trout (*Oncorhynchus mykiss*) was predicted 3.6 times lower affinity (7.6nm) than hACE2 (2.1nm), indicating that Chinook salmon is not susceptible to SARS-CoV-2 infection. These findings suggested frequent detections of SARS-CoV-2 virus in Chinook salmon and even other non-mammal vertebrates might be resulted from unknown contaminations.

Consistent with the findings from non-mammal species, we found consistent affinity phenotype divergences but an expansion of high-affinity phenotypes in 127 mammals, offering possible implication that the rapid expansion of mammals might accelerate the evolution of SARS-CoV-2-like CoVs. As expected, high-affinity phenotypes were significantly enriched in Artiodactyla and Carnivora. Among 48 mammals’ species that were susceptible to infection of SARS-CoV-2 reported by *in vitro* as well as animal infection studies[1, 3, 6, 17-25], 28 animal species were from Artiodactyla and Carnivora orders. In Carnivora, whether SARS-CoV-2 can infect dogs or not triggered some controversies. Shi et al. study found dogs showed low susceptibility to virus and poorly infected[3] but another study considered dogs as intermediate hosts for SARS-CoV-2 virus transmission[24]. Our predicted affinity of dog (*Canis lupus*) (1.6nM) tended to support SARS-CoV-2 infection to dog. Two dogs from households with confirmed human cases of COVID-19 in Hong Kong were found to be infected with SARS-CoV-2, further suggesting that these are instances of human-to-animal transmission of SARS-CoV-2[29]. It is unclear whether infected dogs can transmit the virus to other animals or back to humans. Bosco-Lauth *et al* study suggested that while neither dog nor cat developed clinical disease with the infection of SARS-CoV-2, cats shed infectious virus for up to 5 d and infected naive cats via direct contact, while dogs do not appear to shed virus. Cats that were reinfected with SARS-CoV-2 mounted an effective immune response and did not become reinfected[30]. Pig (*Sus scrofa*) from Artiodactyla order was another controversial animal. Some studies reported pigs were not susceptible to SARS-CoV-2[3, 28], yet other studies reported pig ACE2 could efficiently facilitated virus entry[1, 24]. We found that pig showed a slightly lower affinity (2.3nM) than hACE2 and thought that the infection risk of pig could not be ignored. By contrast, low affinity phenotypes were dominant in Rodentia and Chiroptera. Our predicted phenotypes of rodent animals were consistent with failure cases of SARS-CoV-2 infection in common rat and mouse model[1, 23, 24, 31]. Extreme low affinities were found in most of species in Chiroptera order in which *Rhinolophus sinicus* with the extremely low affinity (68nM) was considered as the natural host of SARS-CoV-2[1]. Extremely low affinity of Chiroptera order species might explain why bats are considered as natural reservoirs of SARS-CoV-like viruses[32, 33]. SARS-Cov-2 infection barely succeeded or succeeded just at very low level in *Rhinolophus sinicus* cells[1, 34, 35]. Nevertheless, SARS-CoV-2 can successfully infect *Rhinolophus sinicus* bat intestinal epithelium organoid[31]. Such differences of infection phenotypes might be partially due to technological bias of the intestinal epithelium organoid simulating real environment of *Rhinolophus sinicus* bat intestines.

Primate animals is the most convincing animal model for evaluating potential drugs and vaccines during the COVID-19 outbreak. In primate orders, we observed slightly high or consistent affinity phenotypes with hACE2 exited in OWMs, with one exception of Golden snub-nosed monkey (2.6nM) with slightly lower affinity. The low-affinity phenotypes occur in NWMs, Tarsiiformes, Lorisiformes as well as Chinese tree shrew (2.6nM) (Scandentia order as the closely relative of Primates). Consistent with our predicted affinity phenotypes, *Macaca mulatta* (1.8nm) and *Macaca fascicularis* (1.9nM) of OWMs were successfully infected by SARS-CoV-2 while Callithrix jacchus (6.8nM) of NWMs failed in SARS-CoV-2 infection[5]. Ocular conjunctival inoculation of SARS-CoV-2 can cause mild COVID-19 in rhesus macaques (*Macaca mulatta*) and could not be re-infected after symptoms were alleviated with the specific antibody tested positively[5, 36]. Cynomolgus macaques (*Macaca fascicularis*) could shed virus for a prolonged period of time with COVID-19-like symptom[25]. These findings suggested that rhesus and cyophagous macaques are appropriate as animal models for evaluating vaccines and drugs for the treatment or prevention of COVID-19.

To determine what amino acid variations of ACE2 contribute to diversified affinity phenotypes is vital for the development of both drug and vaccine during the progress of COVID-19. We found 22 known amino acid residues in ACE2 were unable to explain affinity phenotype diversities from 247 vertebrates. In contrast, at the 8 amino acid sites around 19-82 amino acid region (referred to as hACE2), we found several novel natural mutations contributing to various binding affinity phenotypes. For example, N27/Q27, Q30, Q79 and T82 could increase the hACE2 binding affinity; yet, L19, N34, M79, H41, and deletion of Q24 enable clearly lower the affinity (**Figure 4**). H41 always existed in the extreme low affinity hosts, it could clearly lower the affinity of hosts that was with a higher than hACE2 affinity (**Figure 3c**) if the amino acid change of Y41 to H41 occurs. Structure analyses showed the losing of hydrogen bond to both N501 and T500 of S protein which was formed by Y41, resulting the lower affinity (**Figure 4**). Combining the amino acid residues with the extreme affinity phenotype at each of the 8 sites, we further identified a high-affinity-associated concomitant mutation group (19N(P)-27N-30Q-41Y-79Q-82T) and low-affinity-associated concomitant mutation group (19L-24Del-27A-34N-41H-79M-82M) which could clearly reverse affinity phenotypes between high-or-low-affinity animal species. The hrsACE2 can significantly block early stages of SARS-CoV-2 infections[37]. The combined amino acid residues contributing to extreme higher affinity may provide novel potentials for the development of potential human recombinant soluble ACE2 (hrsACE2) in treating patients with COVID-19 or for constructing genetically modified SARS-CoV-2 infection models promoting vaccines studies.

SARS-CoV-2 uses ACE2 as recognized receptor for its entry into host cell and the virus surface S protein mediates SARS-CoV-2 entry into cells[9, 38, 39], which comprises two functional subunits responsible for binding to the host cell receptor (S1 subunit) and fusion of the viral and cellular membranes (S2 subunit). During SARS-CoV-2 infection, once the RBD of S1 subunit binds to hosts ACE2, S protein is cleaved by host proteases into S1 and S2 subunits at the S2’ site before extensive irreversible conformational changes for the membrane fusion[39, 40]. This cleavage can activate the membrane fusion[39, 40]. Some studies have confirmed that the cathepsin B and L (CatB/L) and TMPRSS2 play important roles in S protein cleavage of SARS-CoV-2[9, 41-43]. B0AT1 (SLC6A19) often serves as a transporter for ACE2 and the presence of B0AT1 may block TMPRSS2 to the cutting site on ACE2[11]. However, whether B0AT1 can suppress SARS-CoV-2 infection by blocking ACE2 cleavage still remain to be explored. Distinct from SARS-CoV, SARS-CoV-2 virus shares a similar furin cleavage sites at S1/S2 sites with MERS-CoV virus[2]. Like MERS-CoV, pre-cleavage at the S1/S2 site mediated by furin protein might promote subsequent TMPRSS2-dependent entry of SARS-CoV-2[39, 44, 45]. Besides ACE2, CD147-SP also could be recruited by SARS-CoV-2 for invading host cells[46]. Using human furin (GeneBank accession: NP_001276752.1), cathepsin L (GeneBank accession: NP_001903), TMPRSS2 (GeneBank accession: NP_005647.3), and CD147-SP (GeneBank accession: BAC76828.1) proteins blasting against Non-redundant protein sequences (NR) in NCBI, we found 308 placental mammals share amino acid sequence identity of furin higher than 90% with human. Only 26 species from *Simiiformes* infraorder share higher than 90% identity of cathepsin L with human and 39 species from *Hominoidea* superfamily share higher than 90% identity with human TMPRSS2. The human CD147-SP protein only shared higher than 90% identity with those of three Apes species, including Sumatran orangutan, chimpanzee, and western lowland gorilla. These results suggested that furin protein is highly conserved but TMPRSS2 and CD147-SP are greatly diverged or lineage specific across vertebrates. Recently, a new host factor Neuropilin-1 was reported associated with SARS-Cov-2 infection [47]. By blasting against NR database, human Neuropilin-1 (GeneBank accession: AAP80144.1) were found shared more than 93% identity with placental mammals. Like furin protein, Neuropilin-1 is also highly conserved. It further indicated that TMPRSS2 and CD147-SP with highly species specificity might contribute to various infection phenotypes to SARS-CoV-2 to different animal hosts. Despite binding to ACE2 is well-known to a critical step for cell entry of SARS-CoV-2 or SARS-CoV-2 like virus, our study suggested that only the affinity testing of ACE2 could not completely estimate SARS-Cov-2 infection. In the future, it is crucial to elucidate pathogenetic mechanism of SARS-CoV-2 by considering comprehensive understanding of combined multiple host factors, such as cleavage proteases or novel functional molecules.

This study provides four major findings for better understanding the evolutionary pattern of bonding affinity of ACE2 in 247 jawed vertebrates to the S protein of SARS-CoV-2. First, high-or-low binding affinity phenotype divergence of ACE2 to the S protein of SARS-CoV-2 has appeared in two ancient species of jawed vertebrates, *Scyliorhinus torazame* (low affinity, Chondrichthyes) and *Latimeria chalumnae* (high affinity, Coelacanthimorpha). Second, multiple independent affinity divergence events recur in fishes, amphibians-reptiles, birds, and mammals, which could be explained to great extent by lineage-specific amino acid mutations. Third, high affinity phenotypes go up in mammals, possibly implying the rapid expansion of mammals might accelerate the evolution of coronaviruses. Fourth, we found natural mutations at eight amino acid sites of ACE2 can determine most of phenotype divergences of bonding affinity in 247 vertebrates and resolved structural basis of divergent bonding affinity phenotypes. Moreover, our identified high-affinity or low-affinity-associated concomitant mutation group would offer potential benefits for the treatment and prevention of SARS-CoV-2. In the future, much more attention was needed focusing on the cleavage proteins to obtain a detail and comprehensive description for preventing SARS-CoV-2 infection.

## Materials and Methods

### Obtaining amino acid sequences of ACE2 and the S protein of SARS-CoV-2

Raw ACE2 amino acid (AA) sequences belonging to vertebrates were downloaded from the nr database from NCBI and UniProt database. After manually removing AA sequences with length < 700aa or duplicated in the same host or that labeled by low quality in sequence title, a total of 247 ACE2 AA sequences representing 247 vertebrates were finally kept. SARS-Cov-2 AA sequence were downloaded from GenBank with accession number MN908947.

### Protein structure homology modeling and affinity prediction between wild ACE2 peptidase domain (PD) and the RBD of the S protein of SARS-CoV-2

To test the bonding affinity between SARS-CoV-2 and vertebrate ACE2, we focused on the bonding affinity between ACE2 PD and the RBD of S protein of SARS-CoV-2. We first aligned ACE2 AA sequences of 247 vertebrates including hACE2 using MEGA X[48] with manually corrections using BioEdit v7.2.5 and then the PD regions ranging from 19 to 615 amino acid residues were extracted from all 247 vertebrates referring to hACE2[11]. The ACE2 protein tree of 247 vertebrates was built using MEGA X[48] and annotated with Interactive Tree Of Life (iTOL) v 5.51[49]. The RBD region of SARS-CoV-2 ranging from 318 to 510 amino acid residues was extracted according to the RBD domain of SARS-CoV BJ01[50].

Protein structure homology modeling were performed using SWISS-modeling workspace[51] using all 247 vertebrates’ ACE2 PD AA sequences and RBD AA sequences of SARS-CoV-2 in automated mode.

Affinity prediction were performed on PRODIGY (PROtein binDIng enerGY prediction, https://bianca.science.uu.nl/prodigy) [16] with pdb file generated by SWISS-modeling. Temperature was set to 37°C.

### Protein structure homology modeling and affinity prediction between vertebrate-derived-hACE2 mutants and the RBD of S protein of SARS-Cov-2

Based on known protein contact residues between SARS-CoV-2 RBD and hACE2[12], we obtained 22 protein contact residues including S19, Q24, T27, F28, D30, K31, H34, E35, E37, D38, Y41, Q42, L45, L79, M82, Y83, N330, K353, G354, D355, R357 and R393 in hACE2. Corresponding to hACE2, we extracted from contact 22 amino acid residues from 247 vertebrates’ ACE2 based on the Mega X aligned file using BioEdit v7.2.5.

We next mutated all 22 residues described above from hACE2 to 247 vertebrates to build vertebrate-derived-hACE2 mutants. The 247 vertebrate-derived-hACE2 PD AA sequence and SARS-CoV-2 RBD AA sequence were used to conduct protein structure homology modeling and affinity prediction according to the methods described above.

### Screening potential amino acid sites contributing to the affinity diversity between 247 vertebrates’ ACE2 PD and SARS-CoV-2 RBD

To find amino acid residues that initially determined the diverse affinity between 247 vertebrates’ ACE2 PD and SARS-CoV-2 RBD, we selected three vertebrates with top high affinity (from at least two orders) and another three vertebrates with top low affinity (from at least two orders) in Actinopteri (High: *Takifugu flavidus, Mastacembelus armatus, Pundamilia nyererei*; Low: *Anarrhichthys ocellatus, Xiphophorus maculatus, Poecilia mexicana*), Aves (High: *Manacus vitellinus, Pipra filicauda, Neopelma chrysocephalum*; Low: *Zonotrichia albicollis, Numida Meleagris, Nothoprocta perdicaria*) and Mammals (High: *Physeter catodon, Procyon lotor, Zalophus californianus*; Low: *Rhinolophus pearsonii, Rhinolophus sinicus, Chrysochloris asiatica*) respectively. For six species from Actinopteri class, if the same amino acid residue appears at given amino acid site in two host species with converse affinity phenotypes, such amino acid sites were excluded from consensus amino acid residues determining high-or-low affinity phenotypes. The remaining consensus amino acid changes of ACE2 in at least two of host species with consistent affinity phenotypes were considered as potentially functional amino acid variations contributing to the affinity diversity. The same standards were performed for six animal species from Aves and those from Mammals. We obtained 12, 31, and 32 putative affinity-associated amino acid sites for Actinopteri class, Aves class, and Mammals class, respectively (**Figure 2A**).

To confirm the potentials of putative affinity-associated amino acid variations in each vertebrate class causing affinity changes, we reconstructed amino acid variants by replacing corresponding amino acid residues in both hACE2 and those species with converse affinity phenotypes. The bonding affinities were estimated based on protein structure homology modeling of mutated ACE2 PD AA sequence and SARS-CoV-2 RBD AA sequence as described above (**Figure S2**).

By integrating putative affinity-associated amino acid sites from three vertebrate classes, we obtained a total of 64 sites each of which could differentiate between high- or-low affinity species in at least one vertebrate class. To trace whether amino acid changes at the 64 sites could reverse bonding affinity of two oldest vertebrate species in our studied cohort, *Latimeria chalumnae* (high-affinity) and *Scyliorhinus torazame* (low-affinity), we cross-replaced amino acid residues at corresponding 64 sites of ACE2 in *Latimeria chalumnae* and *Scyliorhinus torazame*. The built mutants were used to perform homology modeling and affinity prediction according to the method described above (**Figure S2**).

### Identifying key amino acid changes or potential co-variants determining bonding affinity of ACE2 and the RBD of S protein of SAR-CoV-2

To identify key amino acid changes contributing to bonding affinity changes from three vertebrate classes above, we employed a step-by-step splicing strategy to construct a series of mutants. For example, based on our obtained 12 putative affinity-associated sites in each of six species of Actinopteri class, we successively sliced from 1st to 2nd,1st to 3rd, 1st to 4th, …, 1st to 12^th^ sites in Actinopteri species and final obtained 72 aa-slicing groups. Next, we replaced corresponding amino acid residues in hACE2 with those in each slicing group from Actinopteri species and performed protein structure homology modeling and affinity prediction of mutated hACE2 and SARS-CoV-2 RBD as described above. Similar slicing was also performed for those putative affinity-associated sites in Aves class and Mammals class as well as two oldest vertebrate species (**Figures S4-S7**).

According to predicted affinity phenotypes following the change of slicing amino residues in species from each vertebrate class (Actinopteri, Aves, and Mammals), we selected amino acid residues at those sites leading to significant affinity changes as key amino acid residues determining bonding affinity and verified by homology modeling and affinity prediction based on hACE2 mutant building. In turn, we grouped multiple amino acid residues causing strong affinity shift to obtain amino acid co-variants contributing to extremely high or low affinity and verified by homology modeling and affinity prediction based on hACE2 mutants. To further confirm the reliability of amino acid co-variants that boost or lower bonding affinity, we would mutate all amino acid residues from co-variants at corresponding amino acid sites in the ACE2 of all vertebrate species with converse affinity phenotypes (**Figure 3B and 3C**). If predicted affinity phenotypes of at least 95% host species were reversed significantly (at least two-fold changes), thus such amino acid co-variants were selected as candidate targets determining affinity phenotypes.

### The Structures alteration for MT hACE2 mutated by functional AA changes relative to WT hACE2

The 3D Complex Structure presentation was performed using PyMOL v2.0[52] with pdb file generated from protein structure homology modeling with SWISS-modeling workspace in automated mode (Figure 4).

### Quantification and Statistical Analysis

Enrichment analysis of hosts with high or low affinity in each class or each mammalian order were performed with Fisher’s exact test, and statistic p-values were corrected with Benjamini-Hochberg (BH) method. Enrichments with *P*_BH_ <0.05 were considered to be significant.

## Declarations

### Availability of data and materials

The dataset used in this study is provided as supplementary material (Tables S1).

This study did not generate code.

### Ethics approval and consent to participate

Not applicable.

## Acknowledgements

This study was supported by the National Key Research and Development Program of China (no. 2018YFC2000500), the Major Science and Technology Project in Yunnan Province of China (no. 202001BB050001), the Second Tibetan Plateau Scientific Expedition and Research (STEP) program (no. 2019QZKK0503), and the Chinese National Natural Science Foundation (no. 31970571 and U2002206).

## Author Contributions

Z.Z. performed project planning, coordination, execution, and facilitation. T.Z., W.Q., and M.Y. performed modeling analysis. W.Q. and L.W. processed data collection and phylogenetic analysis. Z.Z., Z.C., T.Z., and W.Q. prepared the manuscript.

## Supplemental Information

**Figure S1.**
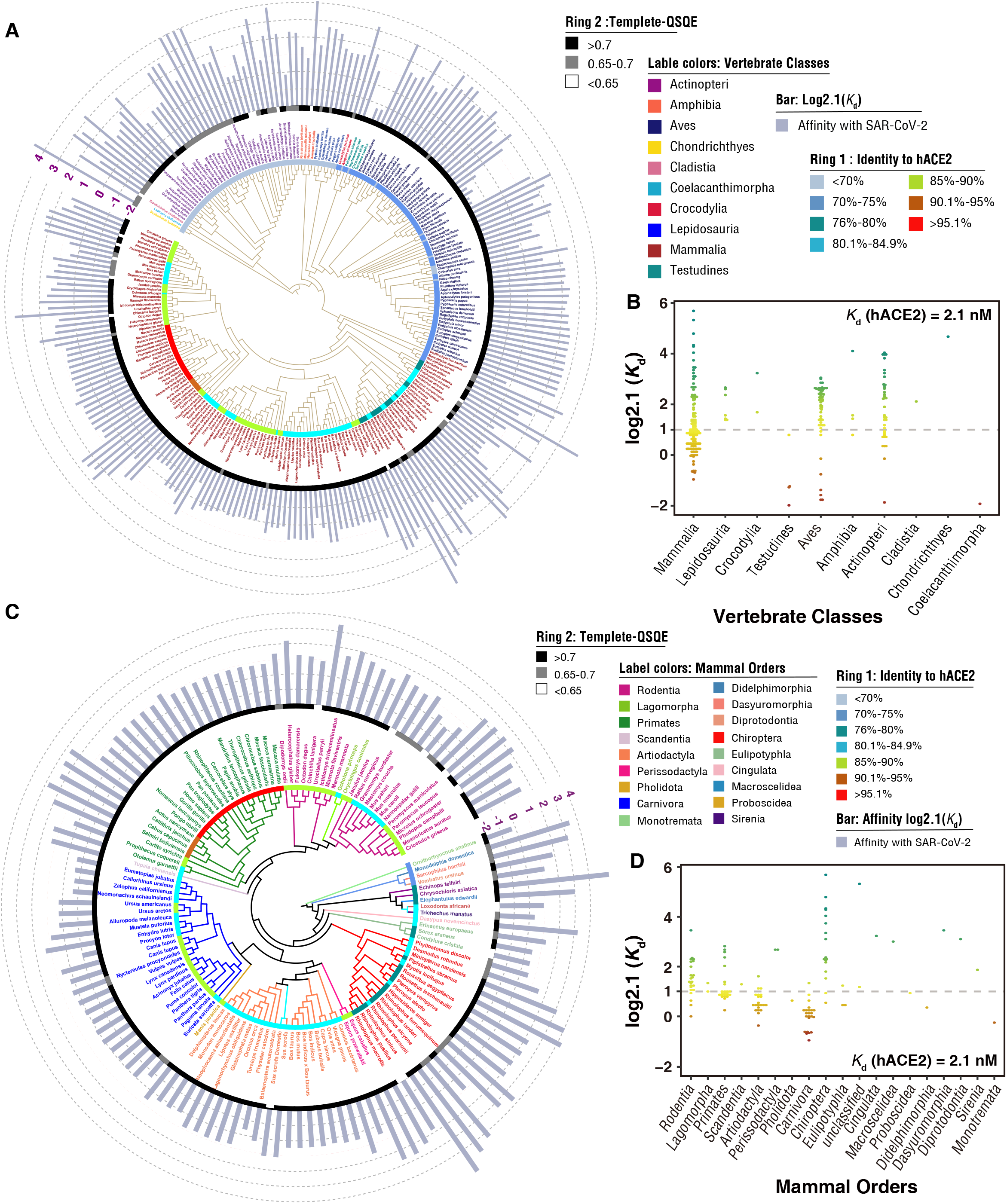
*K*_d_ value details across 247 vertebrates. Linked to Figure 1. **A**. ACE2 protein tree of 247 vertebrates with *K*_d_ value. Bar height indicated value of log2.1(*K*_d_). **B**. *K*d distribution of all host species in each different class of 247 vertebrate hosts. **C**. ACE2 protein tree of 127 mammals with *K*d value. Bar height indicated value of log2.1(Kd). **D**. *K*_d_ distribution of all host species in each order of mammals.

**Figure S2.**
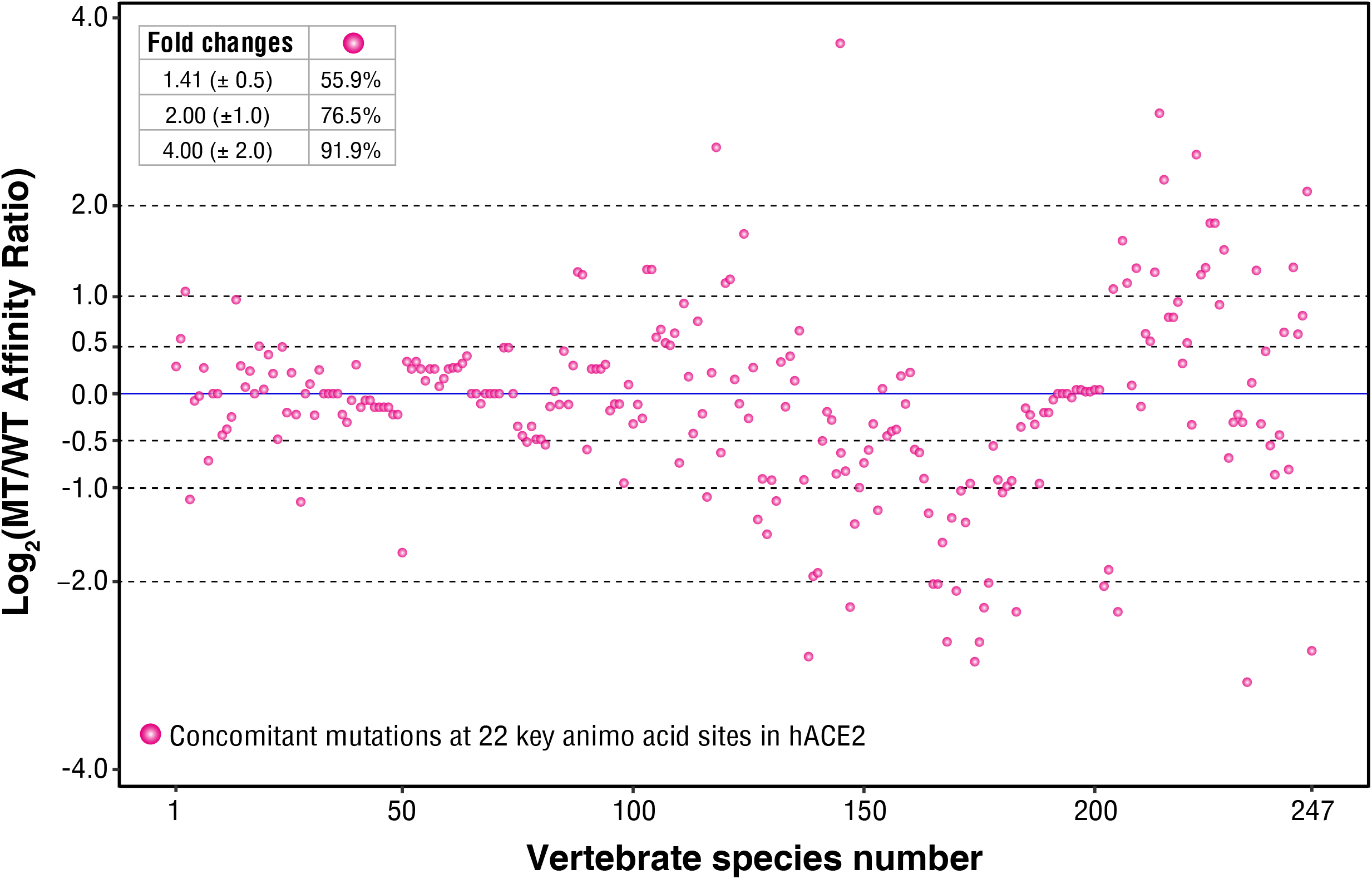
Relative to wild-type (WT) hACE2, affinity changes of mutated (MT) hACE2 based on co-occurring amino acid changes of 247 vertebrates at 22 known contact amino acid sites[12]. Affinity changes were normalized using log2.

**Figure S3.**
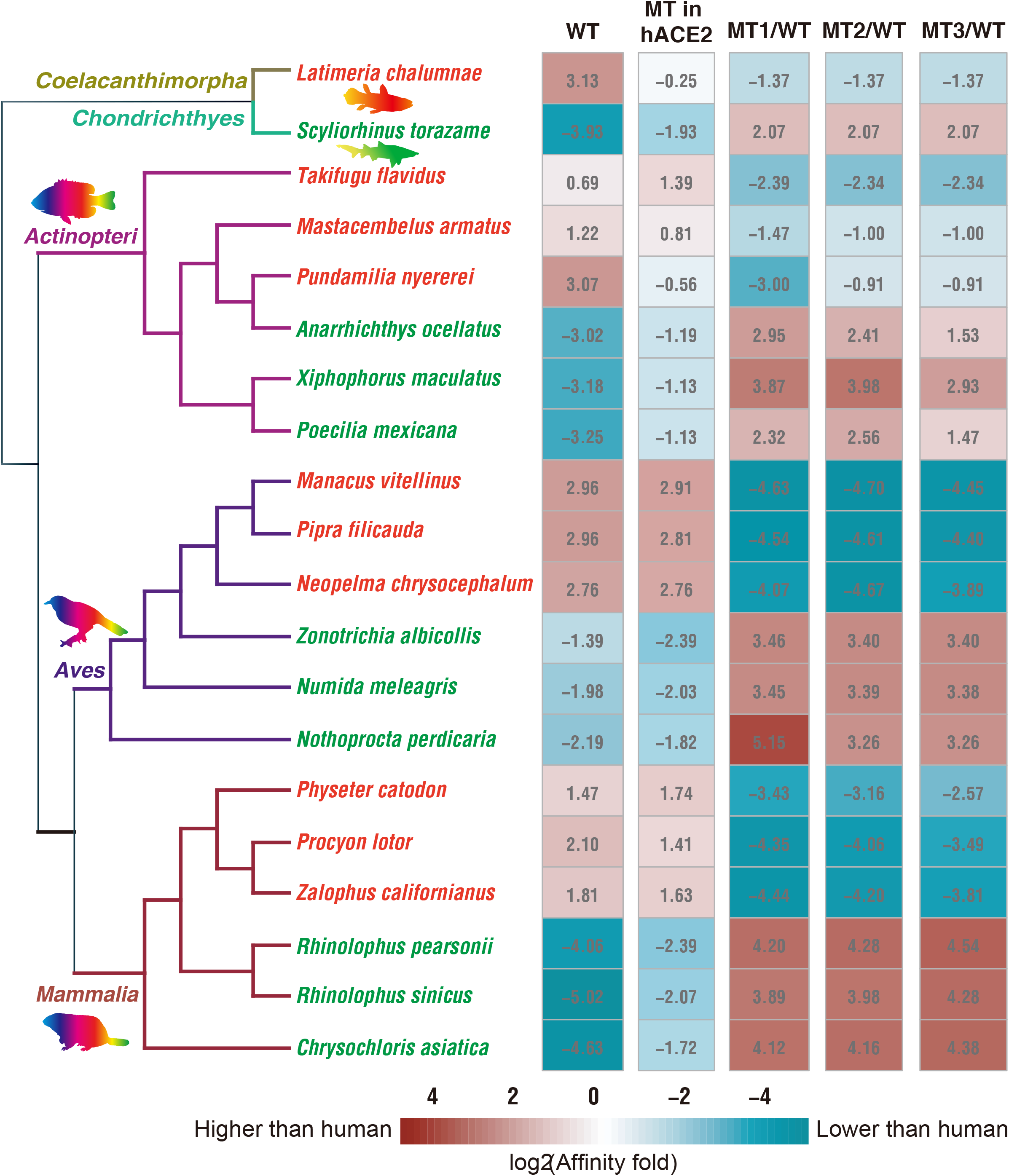
Affinity phenotype reversal after crossed replacements between high- or-low affinity animal species and those animal species in the same lineage with converse phenotypes. Linked to Figure 2. **WT** means affinity fold change of WT ACE2 relative to wild type (WT) hACE2. As controls, **MT in hACE2** means affinity change of mutated (MT) hACE2 using affinity-associated amino acid residues from four animal lineages relative to WT hACE2. Relative to the affinity of WT ACE2 of a given animal species, **MT1-MT3/WT** means affinity change of ACE2 of the animal species replaced by affinity-associated amino acid residues from the first animal species with converse phenotype in same class.

**Figure S4.**
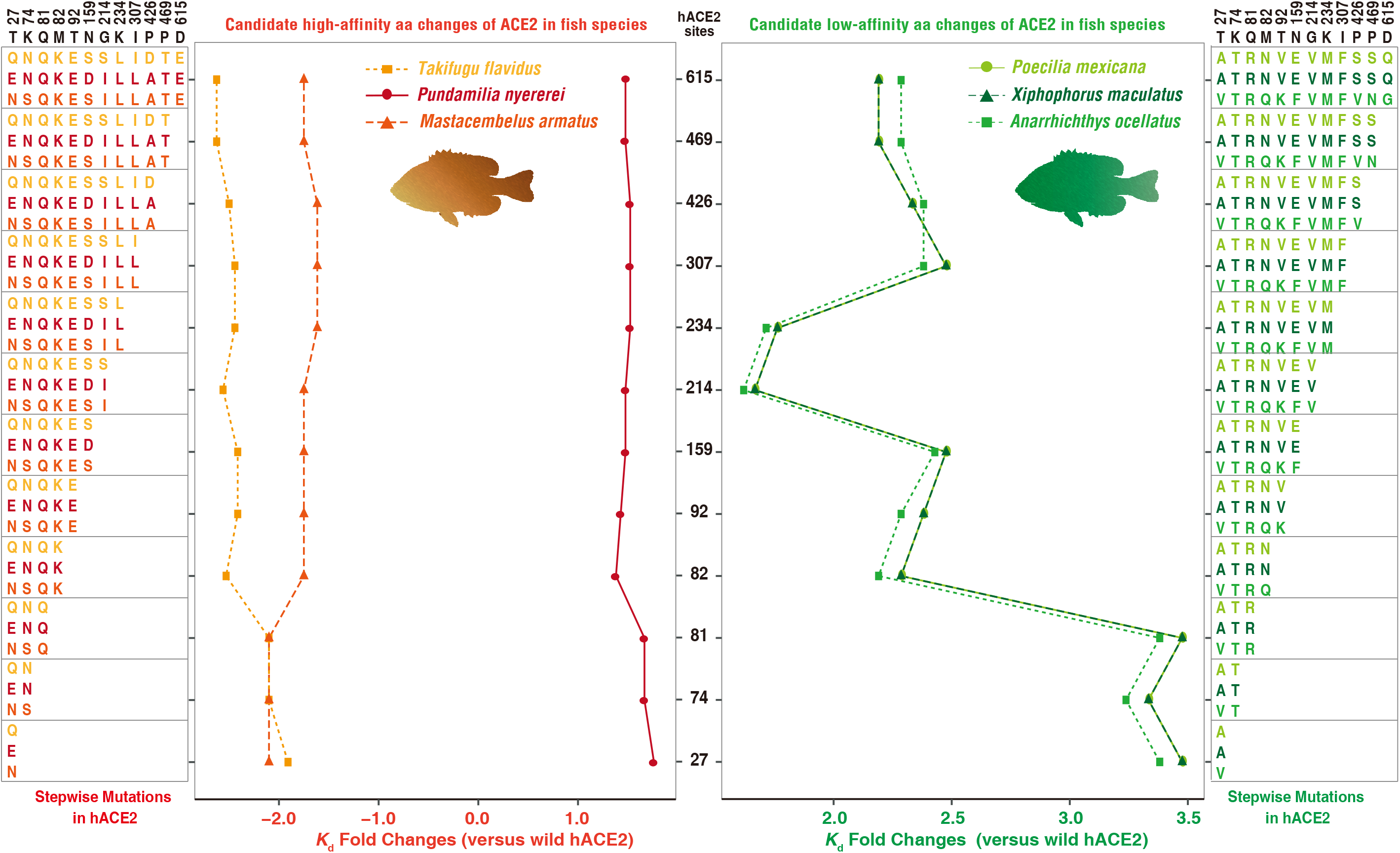
MT-hACE2 affinity changes relative to WT hACE2 following step by step replacement in hACE2 with amino acids at affinity-associated sites from top 3 high affinity hosts (left panel) and top 3 low affinity hosts (right panel) in Actinopteri. Linked to Figures 2 and 3.

**Figure S5.**
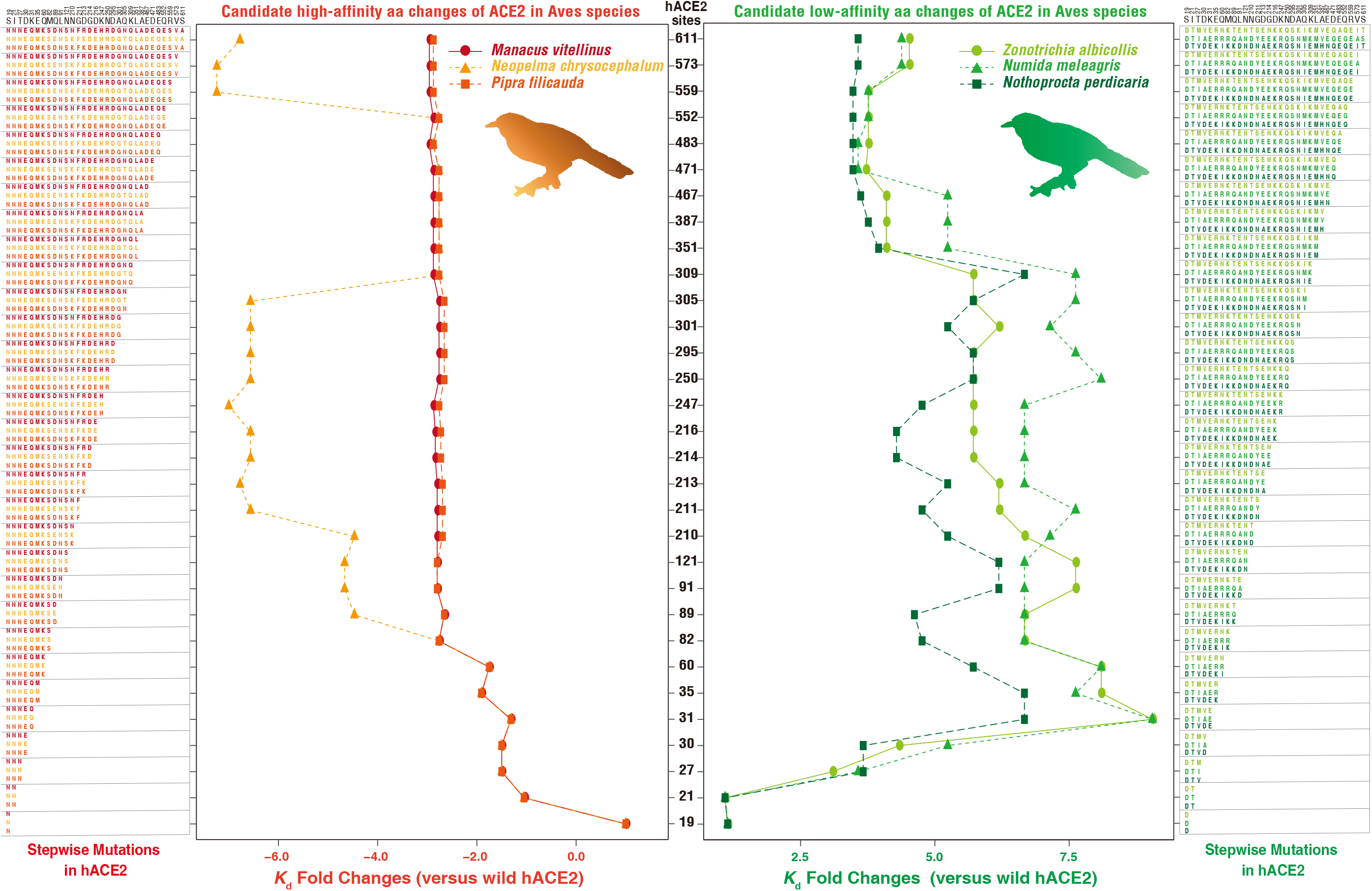
MT-hACE2 affinity changes relative to WT hACE2 following step by step replacement in hACE2 with amino acids at affinity-associated sites from top 3 high affinity hosts (left panel) and top 3 low affinity hosts (right panel) in Aves.

**Figure S6.**
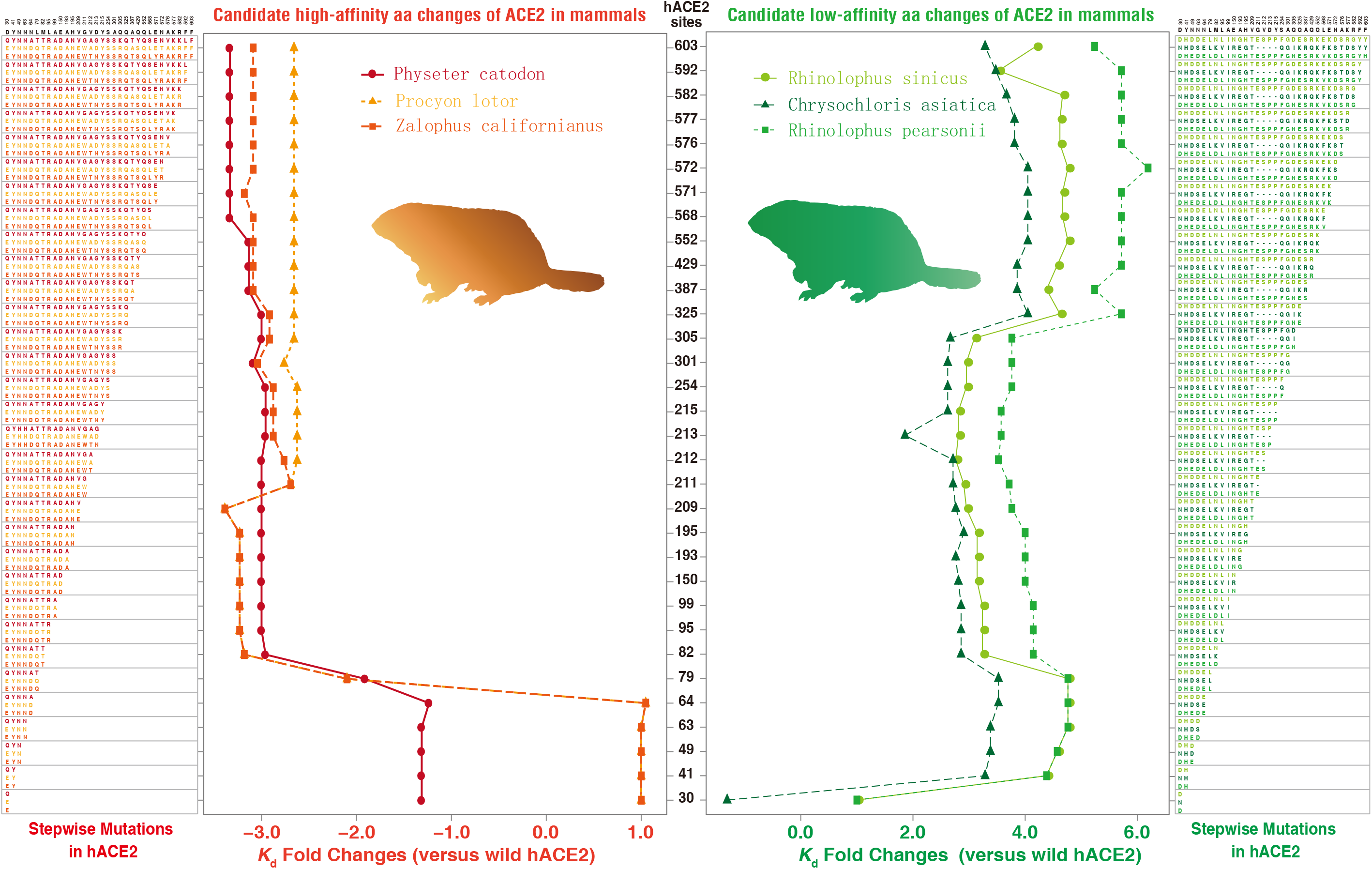
MT-hACE2 affinity changes relative to WT hACE2 following step by step replacement in hACE2 with amino acids at affinity-associated sites from top 3 high affinity hosts (left panel) and top 3 low affinity hosts (right panel) in mammals. Linked to Figures 2 and 3.

**Figure S7.**
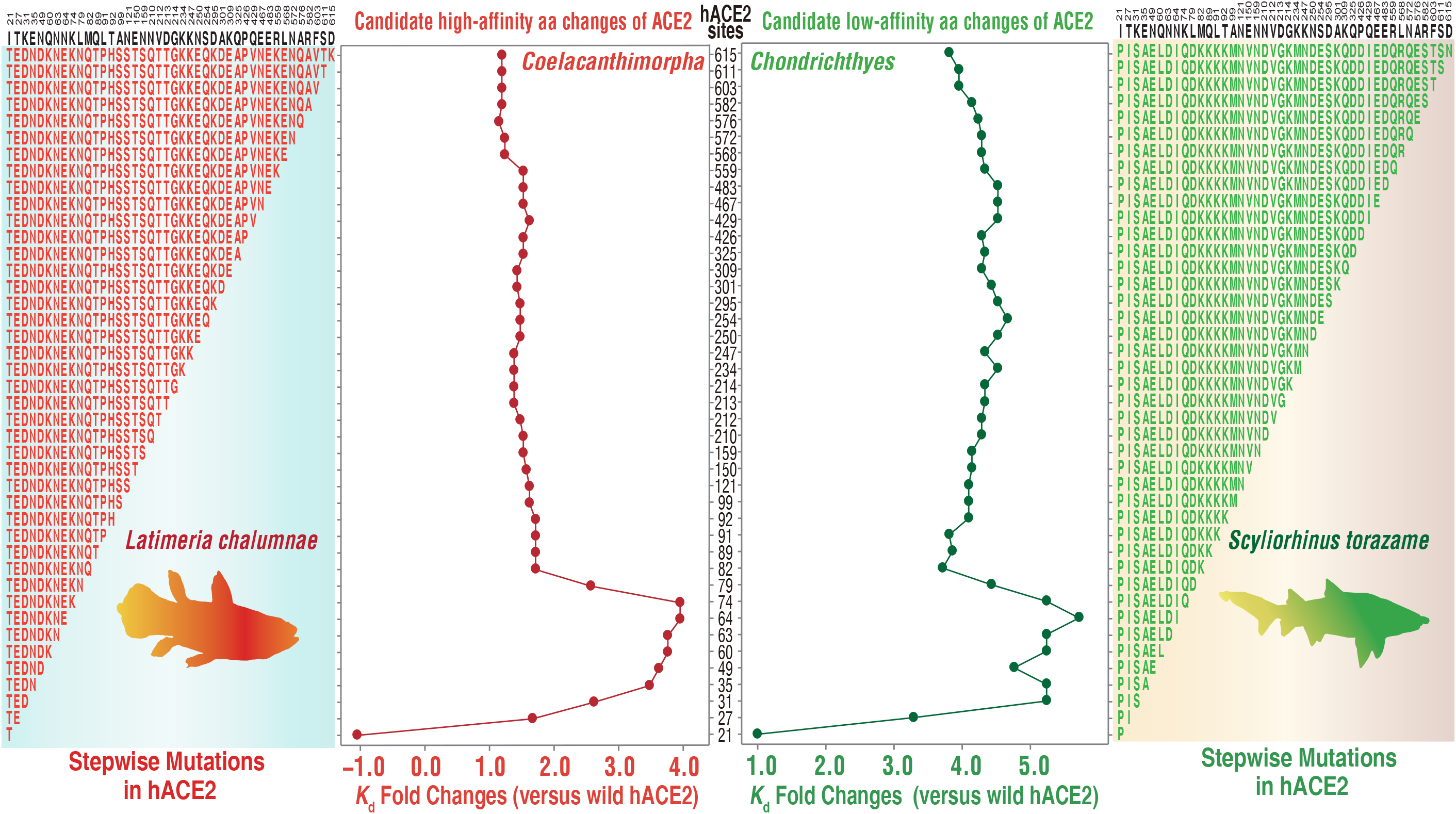
In two ancient jaw vertebrates (left panel: Coelacanthimorpha; right panel: Chondrichthyes), MT-hACE2 affinity changes relative to WT hACE2 following step by step replacement in hACE2 with amino acids linked to affinity-associated sites based on the integration of Actinopteri, Aves, and Mammalian classes in Figures S4-S6. Linked to Figures 2 and 3.

**Figure S8.**
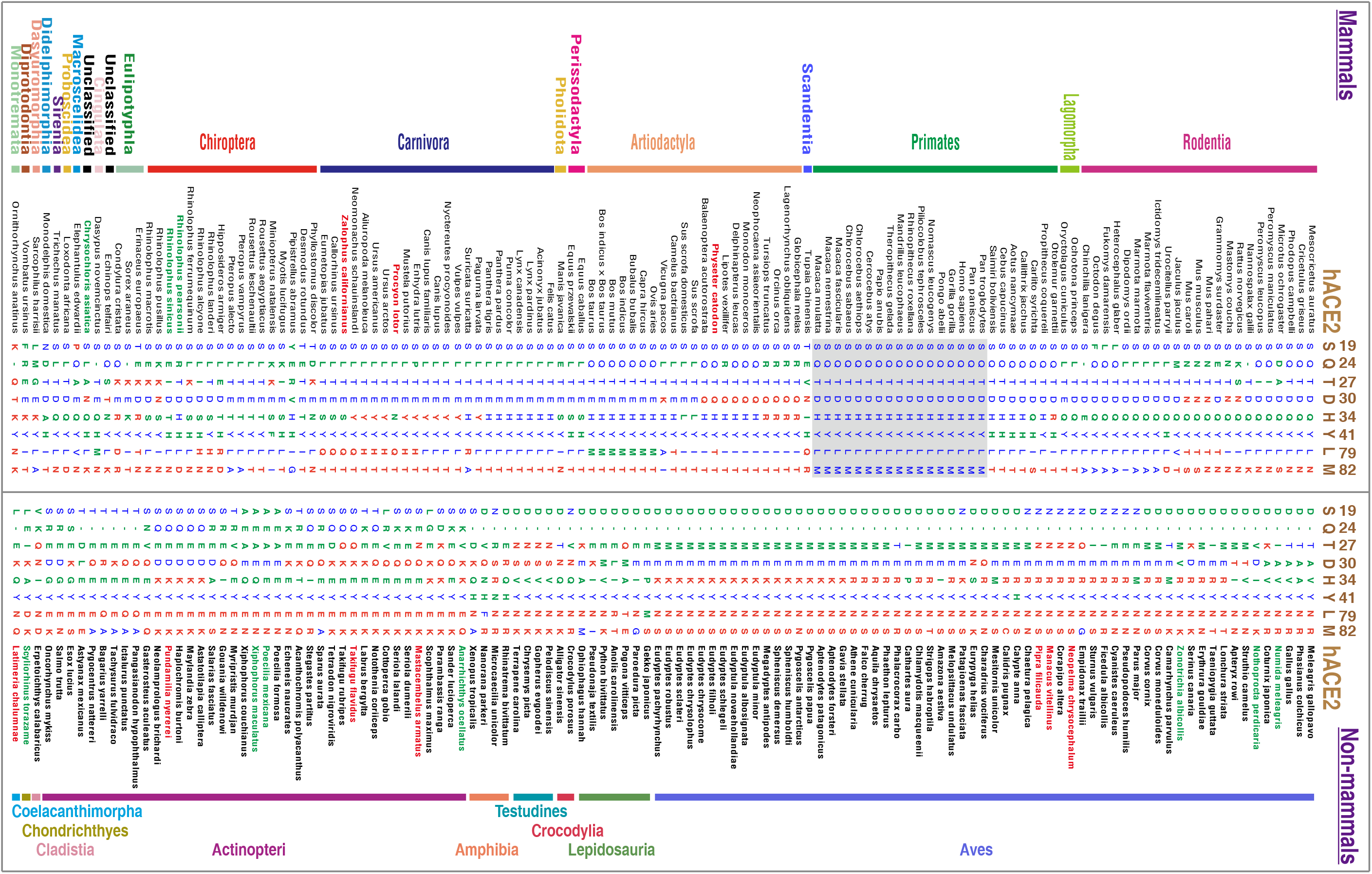
Distribution of amino acid variations at 8 conserved loci across 247 jawed vertebrates. Linked to Figures 2 and 3.

**Table S1. Details of affinity prediction between ACE2 PD from 247 vertebrates and the RBD of the S protein of SARS-COV-2. Linked to Figure 1**.

## References

[1] Zhou P, Yang X-L, Wang X-G, Hu B, Zhang L, Zhang W, et al. A pneumonia outbreak associated with a new coronavirus of probable bat origin. Nature 2020;579:270–3.

[2] Zhang T, Wu Q, Zhang Z. Probable Pangolin Origin of SARS-CoV-2 Associated with the COVID-19 Outbreak. Current Biology 2020;30:1346-51.e2.

[3] Lan J, Ge J, Yu J, Shan S, Zhou H, Fan S, et al. Structure of the SARS-CoV-2 spike receptor-binding domain bound to the ACE2 receptor. Nature 2020.

[4] Oude Munnink BB, Sikkema RS, Nieuwenhuijse DF, Molenaar RJ, Munger E, Molenkamp R, et al. Transmission of SARS-CoV-2 on mink farms between humans and mink and back to humans. Science 2020:eabe5901.

[5] Lu S, Zhao Y, Yu W, Yang Y, Gao J, Wang J, et al. Comparison of nonhuman primates identified the suitable model for COVID-19. Signal Transduction and Targeted Therapy 2020;5:157.

[6] Chan JF-W, Zhang AJ, Yuan S, Poon VK-M, Chan CC-S, Lee AC-Y, et al. Simulation of the Clinical and Pathological Manifestations of Coronavirus Disease 2019 (COVID-19) in a Golden Syrian Hamster Model: Implications for Disease Pathogenesis and Transmissibility. Clinical Infectious Diseases 2020.

[7] Damas J, Hughes GM, Keough KC, Painter CA, Persky NS, Corbo M, et al. Broad host range of SARS-CoV-2 predicted by comparative and structural analysis of ACE2 in vertebrates. Proceedings of the National Academy of Sciences 2020;117:22311–22.

[8] Procko E. The sequence of human ACE2 is suboptimal for binding the S spike protein of SARS coronavirus 2. bioRxiv 2020:2020.03.16.994236.

[9] Hoffmann M, Kleine-Weber H, Schroeder S, Krüger N, Herrler T, Erichsen S, et al. SARS-CoV-2 Cell Entry Depends on ACE2 and TMPRSS2 and Is Blocked by a Clinically Proven Protease Inhibitor. Cell 2020;181:271-80.e8.

[10] Wrapp D, Wang N, Corbett KS, Goldsmith JA, Hsieh C-L, Abiona O, et al. Cryo-EM structure of the 2019-nCoV spike in the prefusion conformation. Science 2020;367:1260–3.

[11] Boehme M, van de Wouw M, Bastiaanssen TFS, Olavarría-Ramírez L, Lyons K, Fouhy F, et al. Mid-life microbiota crises: middle age is associated with pervasive neuroimmune alterations that are reversed by targeting the gut microbiome. Mol Psychiatry 2020;25:2567–83.

[12] Shang J, Ye G, Shi K, Wan Y, Luo C, Aihara H, et al. Structural basis of receptor recognition by SARS-CoV-2. Nature 2020;581:221–4.

[13] Benetti E, Tita R, Spiga O, Ciolfi A, Birolo G, Bruselles A, et al. ACE2 gene variants may underlie interindividual variability and susceptibility to COVID-19 in the Italian population. European Journal of Human Genetics 2020;28:1602–14.

[14] Seo M-H, Park J, Kim E, Hohng S, Kim H-S. Protein conformational dynamics dictate the binding affinity for a ligand. Nature Communications 2014;5:3724.

[15] Vangone A, AMJJ Bonvin. Contacts-based prediction of binding affinity in protein–protein complexes. eLife 2015;4:e07454.

[16] Xue LC, Rodrigues JP, Kastritis PL, Bonvin AM, Vangone A. PRODIGY: a web server for predicting the binding affinity of protein–protein complexes. Bioinformatics 2016;32:3676–8.

[17] Gao Q, Bao L, Mao H, Wang L, Xu K, Yang M, et al. Development of an inactivated vaccine candidate for SARS-CoV-2. Science 2020:eabc1932.

[18] Lam TT, Jia N, Zhang YW, Shum MH, Jiang JF, Zhu HC, et al. Identifying SARS-CoV-2-related coronaviruses in Malayan pangolins. Nature 2020;583:282–5.

[19] Shan C, Yao YF, Yang XL, Zhou YW, Gao G, Peng Y, et al. Infection with novel coronavirus (SARS-CoV-2) causes pneumonia in Rhesus macaques. Cell Res 2020.

[20] Bao L, Deng W, Gao H, Xiao C, Liu J, Xue J, et al. Reinfection could not occur in SARS-CoV-2 infected rhesus macaques. bioRxiv 2020:2020.03.13.990226.

[21] Munster VJ, Feldmann F, Williamson BN, van Doremalen N, Pérez-Pérez L, Schulz J, et al. Respiratory disease and virus shedding in rhesus macaques inoculated with SARS-CoV-2. bioRxiv 2020:2020.03.21.001628.

[22] Kim YI, Kim SG, Kim SM, Kim EH, Park SJ, Yu KM, et al. Infection and Rapid Transmission of SARS-CoV-2 in Ferrets. Cell Host Microbe 2020;27:704–9 e2.

[23] Zhao X, Chen D, Szabla R, Zheng M, Li G, Du P, et al. Broad and Differential Animal Angiotensin-Converting Enzyme 2 Receptor Usage by SARS-CoV-2. Journal of Virology 2020;94:e00940–20.

[24] Liu Y, Hu G, Wang Y, Zhao X, Ji F, Ren W, et al. Functional and Genetic Analysis of Viral Receptor ACE2 Orthologs Reveals Broad Potential Host Range of SARS-CoV-2. bioRxiv 2020:2020.04.22.046565.

[25] Rockx B, Kuiken T, Herfst S, Bestebroer T, Lamers MM, Oude Munnink BB, et al. Comparative pathogenesis of COVID-19, MERS, and SARS in a nonhuman primate model. Science (New York, N.Y.) 2020;368:1012–5.

[26] Lu G, Wang Q, Gao GF. Bat-to-human: spike features determining ‘host jump’ of coronaviruses SARS-CoV, MERS-CoV, and beyond. Trends in Microbiology 2015;23:468–78.

[27] Liu Z, Xiao X, Wei X, Li J, Yang J, Tan H, et al. Composition and divergence of coronavirus spike proteins and host ACE2 receptors predict potential intermediate hosts of SARS-CoV-2. Journal of Medical Virology 2020;92:595–601.

[28] Schlottau K, Rissmann M, Graaf A, Schön J, Sehl J, Wylezich C, et al. SARS-CoV-2 in fruit bats, ferrets, pigs, and chickens: an experimental transmission study. The Lancet Microbe 2020;1:e218–e25.

[29] Sit THC, Brackman CJ, Ip SM, Tam KWS, Law PYT, To EMW, et al. Infection of dogs with SARS-CoV-2. Nature 2020;586:776–8.

[30] Bosco-Lauth AM, Hartwig AE, Porter SM, Gordy PW, Nehring M, Byas AD, et al. Experimental infection of domestic dogs and cats with SARS-CoV-2: Pathogenesis, transmission, and response to reexposure in cats. Proceedings of the National Academy of Sciences 2020;117:26382–8.

[31] Zhou J, Li C, Liu X, Chiu MC, Zhao X, Wang D, et al. Infection of bat and human intestinal organoids by SARS-CoV-2. Nature Medicine 2020.

[32] Lau SKP, Woo PCY, Li KSM, Huang Y, Tsoi H-W, Wong BHL, et al. Severe acute respiratory syndrome coronavirus-like virus in Chinese horseshoe bats. Proceedings of the National Academy of Sciences of the United States of America 2005;102:14040.

[33] Ge X-Y, Li J-L, Yang X-L, Chmura AA, Zhu G, Epstein JH, et al. Isolation and characterization of a bat SARS-like coronavirus that uses the ACE2 receptor. Nature 2013;503:535–8.

[34] Chu H, Chan JF-W, Yuen TT-T, Shuai H, Yuan S, Wang Y, et al. Comparative tropism, replication kinetics, and cell damage profiling of SARS-CoV-2 and SARS-CoV with implications for clinical manifestations, transmissibility, and laboratory studies of COVID-19: an observational study. The Lancet Microbe 2020;1:e14–e23.

[35] Tang Y-D, Li Y-M, Sun J, Zhang H-L, Wang T-Y, Sun M-X, et al. Cell entry of SARS-CoV-2 conferred by angiotensin-converting enzyme 2 (ACE2) of different species. bioRxiv 2020:2020.06.15.153916.

[36] Deng W, Bao L, Gao H, Xiang Z, Qu Y, Song Z, et al. Ocular conjunctival inoculation of SARS-CoV-2 can cause mild COVID-19 in rhesus macaques. Nature Communications 2020;11:4400.

[37] Monteil V, Kwon H, Prado P, Hagelkrüys A, Wimmer RA, Stahl M, et al. Inhibition of SARS-CoV-2 Infections in Engineered Human Tissues Using Clinical-Grade Soluble Human ACE2. Cell 2020;181:905-13.e7.

[38] Li W, Moore MJ, Vasilieva N, Sui J, Wong SK, Berne MA, et al. Angiotensin-converting enzyme 2 is a functional receptor for the SARS coronavirus. Nature 2003;426:450–4.

[39] Walls AC, Park Y-J, Tortorici MA, Wall A, McGuire AT, Veesler D. Structure, Function, and Antigenicity of the SARS-CoV-2 Spike Glycoprotein. Cell 2020;181:281-92.e6.

[40] Tortorici MA, Veesler D. Chapter Four -Structural insights into coronavirus entry. In: Rey F. A. (ed) Advances in Virus Research. Academic Press, 2019, 93–116.

[41] Simmons G, Gosalia DN, Rennekamp AJ, Reeves JD, Diamond SL, Bates P. Inhibitors of cathepsin L prevent severe acute respiratory syndrome coronavirus entry. Proceedings of the National Academy of Sciences of the United States of America 2005;102:11876.

[42] Glowacka I, Bertram S, Müller MA, Allen P, Soilleux E, Pfefferle S, et al. Evidence that TMPRSS2 activates the severe acute respiratory syndrome coronavirus spike protein for membrane fusion and reduces viral control by the humoral immune response. Journal of virology 2011;85:4122–34.

[43] Matsuyama S, Nagata N, Shirato K, Kawase M, Takeda M, Taguchi F. Efficient Activation of the Severe Acute Respiratory Syndrome Coronavirus Spike Protein by the Transmembrane Protease TMPRSS2. Journal of Virology 2010;84:12658.

[44] Kleine-Weber H, Elzayat MT, Hoffmann M, Pöhlmann S. Functional analysis of potential cleavage sites in the MERS-coronavirus spike protein. Scientific Reports 2018;8:16597.

[45] Park J-E, Li K, Barlan A, Fehr AR, Perlman S, McCray PB, Jr., et al. Proteolytic processing of Middle East respiratory syndrome coronavirus spikes expands virus tropism. Proceedings of the National Academy of Sciences of the United States of America 2016;113:12262–7.

[46] Huang C, Wang Y, Li X, Ren L, Zhao J, Hu Y, et al. Clinical features of patients infected with 2019 novel coronavirus in Wuhan, China. The Lancet 2020;395:497–506.

[47] Daly JL, Simonetti B, Klein K, Chen KE, Williamson MK, Antón-Plágaro C, et al. Neuropilin-1 is a host factor for SARS-CoV-2 infection. Science 2020;370:861–5.

[48] Kumar S, Stecher G, Li M, Knyaz C, Tamura K. MEGA X: Molecular Evolutionary Genetics Analysis across Computing Platforms. Molecular Biology and Evolution 2018;35:1547–9.

[49] Letunic I, Bork P. Interactive Tree Of Life (iTOL) v4: recent updates and new developments. Nucleic Acids Research 2019;47:W256–W9.

[50] Li F, Li W, Farzan M, Harrison SC. Structure of SARS Coronavirus Spike Receptor-Binding Domain Complexed with Receptor. Science 2005;309:1864–8.

[51] Bordoli L, Kiefer F, Arnold K, Benkert P, Battey J, Schwede T. Protein structure homology modeling using SWISS-MODEL workspace. Nature Protocols 2009;4:1–13.

[52] Schrodinger, LLC (2015), ‘The PyMOL Molecular Graphics System, Version 1.8’.

